# Circlator: automated circularization of genome assemblies using long sequencing reads

**DOI:** 10.1101/023408

**Authors:** Martin Hunt, Nishadi De Silva, Thomas D Otto, Julian Parkhill, Jacqueline A Keane, Simon R Harris

**Affiliations:** Wellcome Trust Sanger Institute, Wellcome Trust Genome Campus, Cambridge, CB10 1SA, UK

## Abstract

The assembly of DNA sequence data into finished genomes is undergoing a renaissance thanks to emerging technologies producing reads of tens of kilobases. Assembling complete bacterial and small eukaryotic genomes is now possible, but the final step of circularizing sequences remains unsolved. Here we present Circlator, the first tool to automate assembly circularization and produce accurate linear representations of circular sequences. Using Pacific Biosciences and Oxford Nanopore data, Circlator correctly circularized 26 of 27 circularizable sequences, comprising 11 chromosomes and 12 plasmids from bacteria, the apicoplast and mitochondrion of *Plasmodium falciparum* and a human mitochondrion. Circlator is available at http://sanger-pathogens.github.io/circlator/.

## Introduction

The challenge of *de novo* sequence assembly has existed ever since the invention of the first automated DNA sequencers. The assembly of early genome sequence data was largely based on two strategies: BAC/YAC tiling or whole-genome shotgun [1]. Although these strategies allow production of high quality sequences, which are often used as reference genomes today, they are both slow and expensive, and the final stage of completing and, where necessary, circularizing the sequences requires laborious and costly manual finishing. The arrival of high-yielding, short read sequencing technologies drastically reduced the time and cost required to generate high depth whole-genome sequencing data ideal for identification of population variation. However, *de novo* genome assemblies from these data are typically too fragmented for genome completion to be practical, and consequently most short-read assembly algorithms do not tackle issues such as circularization of completed genomes.

The recent availability of high-throughput, long-read (up to tens of kilobases) sequencing technologies, in particular Pacific Biosciences (PacBio) and Oxford Nanopore Technologies (ONT), has again improved the contiguity of automated *de novo* assemblies to the point where production of a single contig per DNA molecule is now possible on an unprecedented scale for bacterial and small eukaryotic genomes [2]. In comparison with short read technologies the per base error rate of PacBio and particularly ONT reads is high, but this has been mitigated by taking advantage of the high yield and random error model of these sequencers. By correcting errors in the raw sequencing reads using self-mapping, high quality long sequences can be produced that are then contiguated using an overlap layout consensus assembly approach (for example, using HGAP [3], PBcR [4] and SPRAI [5]).

Although these new technologies raise the prospect of routine automated completion of genome sequences, current long-read assembly software still typically assumes that the contigs they produce are linear. On the contrary, the genome of almost every species contains at least one circular DNA structure, such as bacterial chromosomes and plasmids and the plastid and mitochondrial genomes of eukaryotes. Correct completion and circularization of these molecules is essential if they are to be used routinely in clinical practice. Whole genome sequencing is already providing improved resolution in bacterial epidemiology [6] and allowing *in silico* prediction of antimicrobial-resistance (AMR) [7]. Many important AMR and virulence determinants are carried on plasmids, illustrating the importance of having complete and accurate information for these circular sequences. Similarly, in humans, the mitochondrial genome has been implicated in controlling phenotypes such as depression [8],[9], Leber hereditary optic neuropathy [10], and myopathy and diabetes mellitus [11].

It is, therefore, clearly important to be able to automatically produce accurate representations of circular DNA structures. However, the linear contigs produced by assembly programs to represent circular DNA structures can contain errors. Near-identical overlaps are often found at each end of the contig, which require significant manual intervention to resolve (Figure 1a). Alternatively, the sequence may be incomplete, with a short sequence that would join the contig ends absent (Figure 1b). Finally, when the sequence being assembled is shorter than the length of some reads, it may contain significant misassemblies in the form of multiple duplications of the entire circular molecule (Figure 1c). Currently, there are two main approaches to resolving the circular structure, based on using BLAST [12] and Minimus2 [13] to identify common sequence at each contig end. In our evaluations these methods fail to address the significant incorrect structural representations produced by *de novo* assemblers and therefore require significant further manual examination to produce correct representations of circular DNA structures.

**Figure 1:**
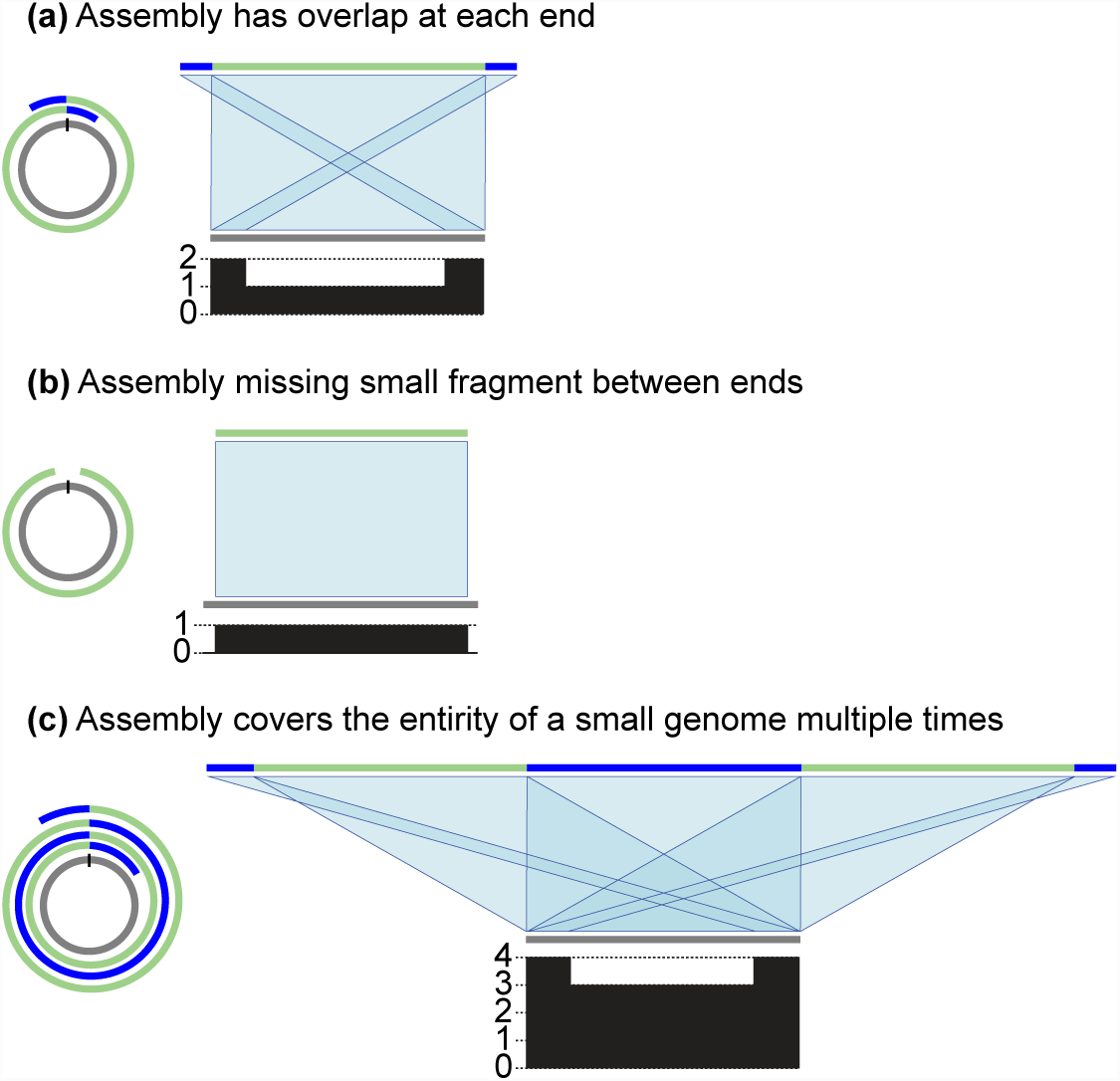
Typical issues in contigs produced by long read assemblers representing circular sequences. In each example, the assembly is in a single contig, coloured with a mix of green and blue, and the reference is shown in gray. Matches between the reference and assembly are shown in light blue. The plot below each reference sequence shows the number of matches to the assembly at each position of the reference sequence. (a) The contig has low quality ends representing the same sequence, which needs resolving into one sequence. (b) The contig has missing sequence. (c) A small circular sequence is assembled into multiple tandem copies.

In this paper we introduce Circlator, a post-assembly improvement toolkit for producing correctly represented circular DNA structures. It uses local assemblies of corrected long reads at contig ends to circularize contigs. This avoids searching for sequence in common between low quality contigs ends, and allows circularization even when overlaps are not present. We evaluated Circlator using examples from a wide range of bacteria, a human genome and a *Plasmodium* sample and show that it outperforms current methods.

## Methods

The Circlator workflow (Supplementary Figure S1) consists of iteratively merging together contigs followed by running local assemblies of corrected reads that align to contig ends. These local assemblies are used to identify circular sequences, each of which is transformed into a linear representation of a circular sequence. Next, the assembly is cleaned by removing small contigs and non-circular contigs that are completely contained within the sequence of another contig. Finally, each circular contig has its start position set to a specified gene (by default dnaA) if present, otherwise the start of a predicted gene near its centre is used.

### Read filtering and local assembly

Circlator takes as input an assembly in FASTA format and the corrected reads that were used to produce that assembly in FASTA or FASTQ format. Both files are produced by common long read assemblers, such as HGAP, PBcR and SPRAI. The corrected reads are mapped to the assembly with BWA-MEM [14] using the -x pacbio option. Reads are then filtered for use in subsequent steps as follows (Supplementary Figure S2). For long contigs (by default, of length at least 100,000bp), only reads that map to the first and last 50,000 bases of the contig are retained.

A read that maps over the position 50,000 bases from either end of the contig is trimmed, so that only the part of the read mapping within 50,000 bases from end of the contig is retained, provided the read is at least 250 bases long after trimming. The remaining reads, which are either mapped to short contigs (*<*100,000bp) or are unmapped, are all retained. These filtered reads are then assembled using SPAdes with the options `--careful --only-assembler` to disable SPAdes’ own correction algorithm and assemble with high stringency. The longest allowed kmer length of 127 is used to maximise the contiguity of the assembly, however SPAdes can fail with this kmer length if the read coverage is low. If SPAdes fails then the kmer length is reduced until an assembly is produced, using values 121, 111, 101, 95, 91, 85, 81, 75 and 71.

### Contig merging

The contigs of the resulting SPAdes assembly are aligned to the original assembly using the nucmer program of MUMmer [16] with options `--maxmatch --diagdiff 25`, and hits with identity of at least 95% are retained using `delta-filter -i 95`. The alignments to each SPAdes contig are analysed to decide if that contig can be used to merge two of the original assembly contigs, as follows (see Figure 2a). The longest match to each of the start and end of the SPAdes contigs is identified. If these matches are to different original assembly contigs, nucmer did not report another longer match to the SPAdes contig, and the positions and orientations are such that the original contigs can be joined, then a new merged contig is constructed. By default, the matches must be at least 4000bp long, within 1000bp of the start or end of the SPAdes contig and within 15000bp of the start or end of the original assembly contigs. More tolerance is allowed in the assembly contigs because they often start and end with low quality sequence. When a join is made, the filtered reads are remapped to the new merged assembly and the process of read filtering, assembly and contig merging is repeated until no more contigs can be merged.

**Figure 2:**
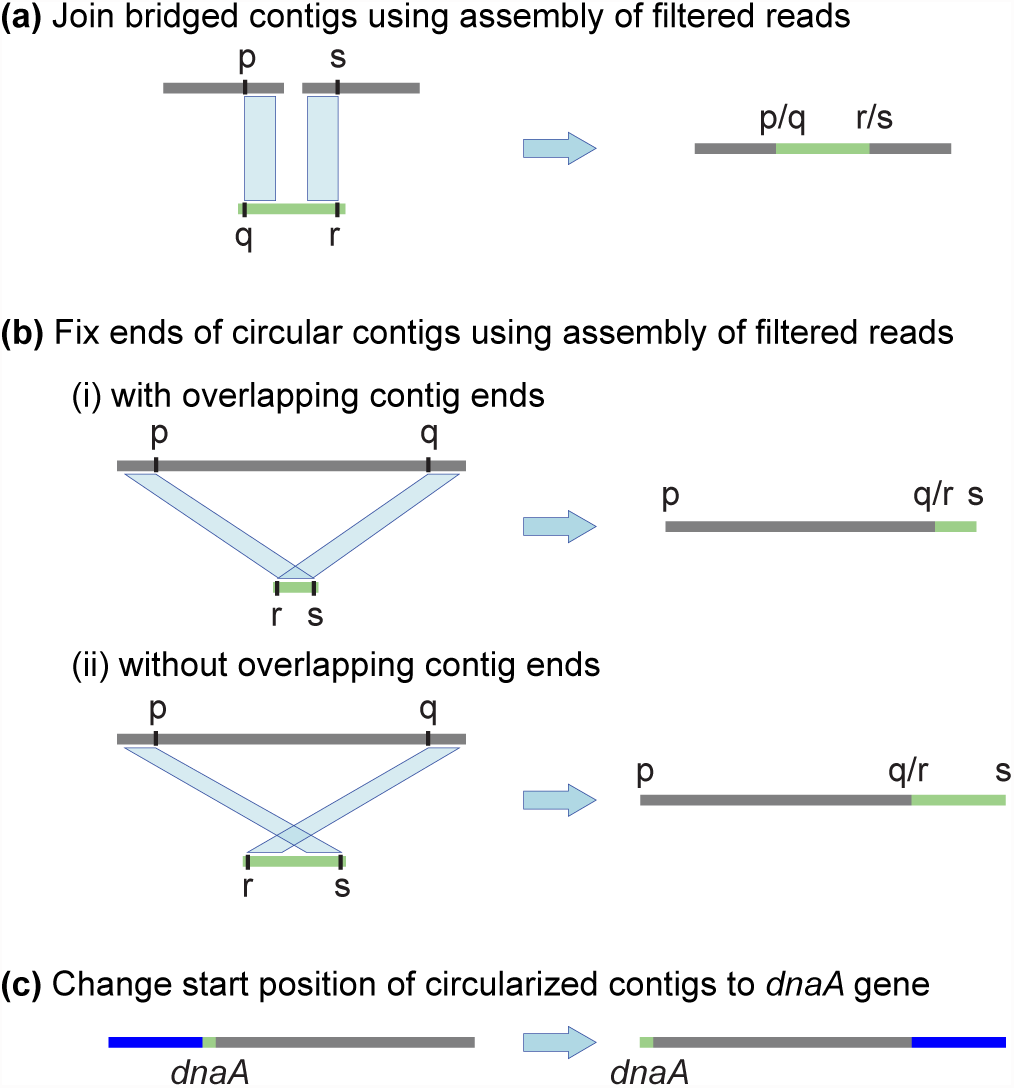
Key stages of the Circlator pipeline. (a) Before circularization, input contigs are merged using *de novo* assemblies of filtered reads. (b) Circular contigs are resolved using matches to contigs assembled from filtered reads. (c) Circularized contigs are rearranged to start at the dnaA gene, or a different gene specified by the user.

### Circularization

Once all possible contig merges are made, the final SPAdes assembly from iterative merging is again aligned to the merged assembly using nucmer with the same settings. Circlator attempts to circularize each contig in turn. First, an attempt is made to match the contig to a SPAdes contig that was identified as circular by SPAdes. If the original assembly contig has nucmer matches that cover at least 95% of its length, all to the same SPAdes circular contig, and one of those nucmer matches has length at least 95% of the length of the SPAdes contig, then the original assembly contig is replaced with the SPAdes contig. If no such SPAdes contig is found, then the longest match to the start and to the end of the original contig is identified, using the same criteria as in the merging stage above. If these two matches are the same, then the second longest match is used. If the two matches are to the same SPAdes contig, and are in the correct orientations and positions (as in Figure 2b), then the contig is circularized. If neither method works, then the original assembly contig is not changed.

### Contig refinement

The assembly is refined further by discarding all contigs shorter than a minimum length, by default 2000bp. Next, all non-circular contigs that are found to be redundant are removed, using the following method. The assembly is aligned to itself using nucmer with the same options as above for contig merging. Contig A is said to be contained in contig B if it has not previously been identified as circular, and there is a nucmer alignment to contig B with at least 95% identity and of length at least 95% of the length of contig A. All such instances of contained contigs are identified, and the relationships are expanded using transitivity. For example, if A is contained in B and B is contained in C, then we ensure that A is contained in C (in case nucmer did not report that A is contained in C). For each set of equivalent contigs, specifically where every contig of the set is contained in all other contigs in the set, only the longest contig of the set is retained. Finally, each remaining contig that is contained in another contig is removed from the assembly.

The last stage of the pipeline rearranges all circular contigs to begin at a known gene. By default each contig is searched against the nucleotide sequences of 162 dnaA genes obtained from RefSeq [17] (see Supplementary Materials for details). The user can provide any alternative FASTA file of genes, but dnaA is used by default because this gene is found in most bacteria and is usually close to the origin of replication [18]. The search is performed on translated nucleotide sequences using the PROmer algorithm of MUMmer, and hits with a minimum percent identity of 80 are retained using `delta-filter -i 80`. The first match to the entire length of any dnaA gene is used to rearrange the contig so that it starts with that gene on its forward strand. If no such match is found, the start of the gene nearest to the middle of the contig is used instead, identified using Prodigal [19] with the option -c to prevent prediction at contig ends.

### Optimising existing methods

In addition to developing Circlator, two existing methods were modified and automated, for comparison with Circlator. The modifications were made in order to improve the effectiveness and accuracy of these methods at circularizing assemblies. We shall refer to these two BLAST- and Minimus2-based methods as simply “BLAST” and “Minimus2” for the remainder of the manuscript.

#### BLAST method

One approach to solving the circularization problem is to use BLAST to identify sequence that is in common at the start and end of a contig. This has been implemented in a script called `check_circularity.pl` included with the SPRAI assembler. Whenever an overlap is found between the start and end of a contig, it is identified as circular and the duplicated sequence at the start of the contig is removed. A circular sequence that is shorter than twice the read length is often assembled into a contig consisting of multiple tandem copies of the original sequence, for example see Figure 1c. In this situation, trimming one duplicated sequence is not sufficient to fix the contig because multiple copies of the true sequence remain. To solve this, we developed a script (`sprai check circularity iterative.py`, included in Supplementary material) that iteratively runs the check circularity.pl script until no more sequence can be removed from any contig ends. The `check circularity.pl` script included in SPRAI version 0.9.9.1 was used, which in turn ran blast+ version 2.2.30 [20].

#### Minimus2 method

The protocol recommended by PacBio to identify and repair circular contigs [21] is based on using Minimus2. A copy of this protocol is included in the Supplementary Material. We automated this manual protocol, with the improvements described below, and added it as an option to Circlator. Minimus2 from version 3.1.0 of the AMOS suite was used. First, Minimus2 is run on the input assembly to merge any overlapping contigs (this is optional, and not part of the original protocol). An attempt is then made to circularize each contig in turn (see Supplementary Figure S3) by breaking it in half to make two smaller contigs, and using the two smaller contigs as input to Minimus2. If the original contig had sequence in common between its start and end, then this should be recognised by Minimus2 and the two smaller contigs will be merged into a single circularized contig. If Minimus2 outputs one contig then this contig is used, otherwise the original, unbroken, contig is kept. The original protocol involved breaking every contig in half and running Minimus2 once on all of the broken contigs. We treat each contig separately because Minimus2 can incorrectly merge parts of different contigs when it is run on all the contig halves pooled together. Moreover, Minimus2 often fails to run, in which case the original contig is retained, whereas the original protocol would produce no output.

### Assembly Polishing

After circularization with any of the three methods implemented here it is important to correct errors in the assembly using raw sequencing reads, since it can contain single base errors and small insertions and deletions. For PacBio data, this polishing step is usually carried out using the Quiver [3] algorithm included in the SMRT-Analysis software package. Nanopolish [22] can be used to correct errors using nanopore data.

## Evaluation

To test the applicability and scalability of Circlator, it was evaluated on 14 bacterial genomes, the circular apicoplast and mitochondrion genomes of the malaria parasite *Plasmodium falciparum*, and the mitochondrion genome of *Homo sapiens*. Finally, we evaluated Circlator on an assembly of the bacterium *Escherichia coli* based on Oxford Nanopore data. In all cases, Circlator was compared against the BLAST and Minimus2 circularization methods described above. Comparisons of all reference sequences and input and output assemblies are shown in Supplementary figures S5-22. Default settings were used for all programs, except where noted for the nanopore and *P. falciparum* data. Circlator version 0.14.0 was used, with SPAdes version 3.5.0, MUMmer version 3.23, BWA-MEM version 0.7.12, Prodigal version 2.60 and SAMtools [23] version 0.1.9.

### Bacterial PacBio data

The evaluated panel of 14 bacterial strains included both gram positive and negative species selected from the National Collection of Type Cultures (NCTC) 3000 project [24] on the basis of there being high-quality reference genome sequences of the same strains available for comparison (see Supplementary Table S1 for sample and reference genome accession numbers). In total, the reference genomes of the sequenced strains comprised 14 chromosomes and 14 plasmids. A per-strain summary of the merging and circularization of the three evaluated methods is shown in Table 1, and a more detailed version is given in Supplementary Table S2.

**Table 1:**
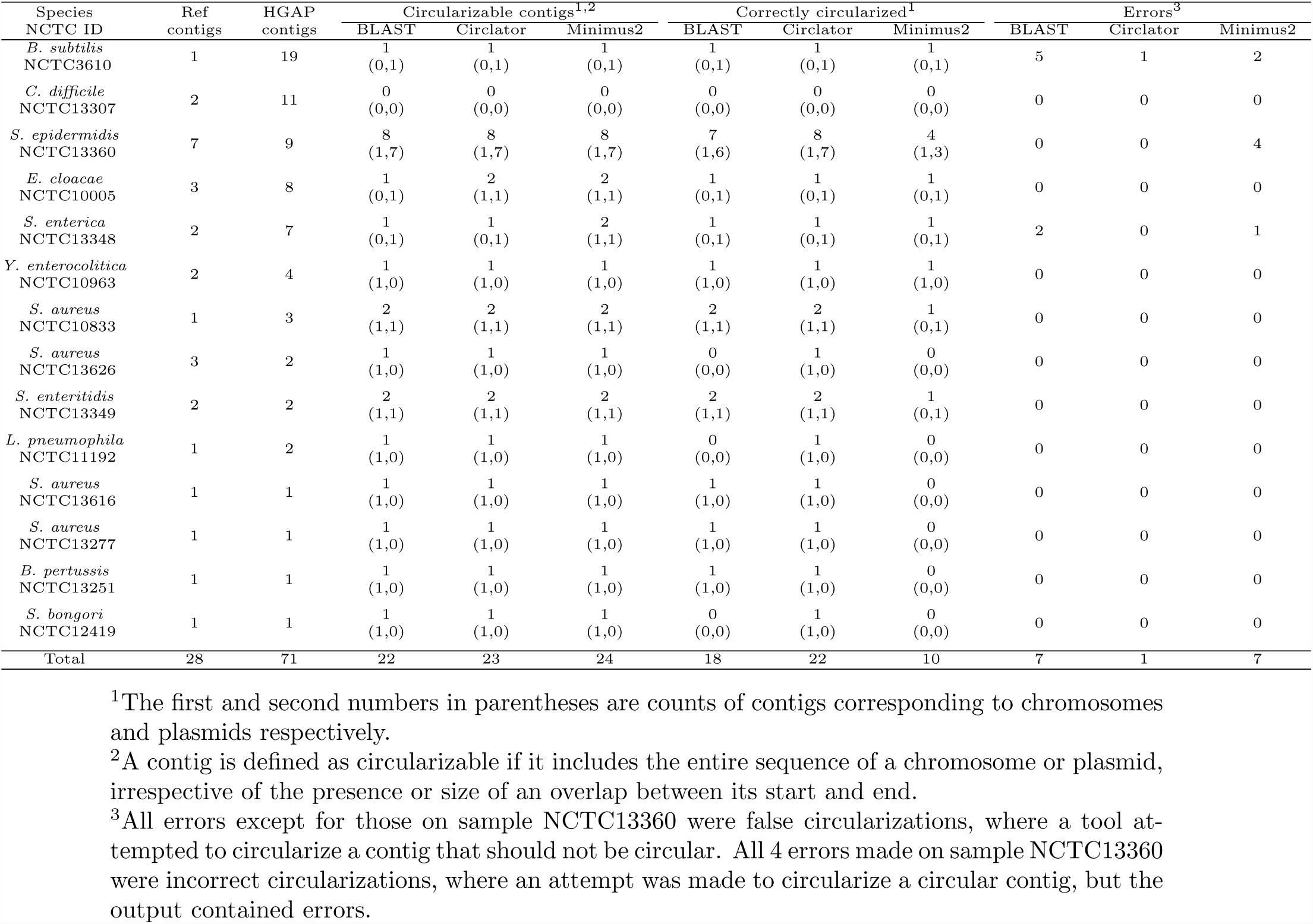
Summary of results on 14 bacterial genome assemblies.

#### Assembly

Assembly of our panel of bacterial genomes using HGAP produced a total of 71 contigs, of which 10 represented complete chromosome sequences and a further 12 represented complete plasmid sequences. Four of these plasmids were not present in the corresponding reference genomes and one plasmid from the reference genome of NCTC13626 was not represented in the HGAP assembly of the same strain. Comparison of the HGAP assemblies to the reference genomes showed that in four cases large regions of the assembled chromosomes were inverted relative to the reference genome. We suspect these represent misassemblies, but assessed that these inversions did not affect the possibility of circularizing the assembled chromosomes.

#### Merging

After merging, Circlator reduced the total number of contigs in the assemblies to 52, while Minimus2 made more merges, reducing the number of contigs to 48. In all cases, manual inspection confirmed the merges to be correct.

#### Circularizable contigs

Contigs were considered circularizable if they included the entire sequence of a chromosome or plasmid, irrespective of the presence or size of an overlap between the two ends of the contig. 22 contigs (10 chromosomes and 12 plasmids), were circularizable under this definition in the initial HGAP assemblies, and as a result of merging contigs, Circlator produced one and Minimus2 two extra circularizable chromosomes. All other chromosomes and plasmids were in multiple contigs, so could not be circularized. Therefore, the number of circularizable contigs for the three methods were 23 (11 chromosomes and 12 plasmids) for Circlator, 24 (12 chromosomes and 12 plasmids) for Minimus2 and 22 (10 chromosomes and 12 plasmids) for BLAST.

#### Circularization

Circlator correctly circularized 22 of 23 circularizable contigs. In comparison, Minimus2 correctly circularized 10/24, and BLAST 18/22. Minimus2 performed particularly badly at circularizing chromosomal contigs, for which it succeeded in only 2 of 12 cases.

Where assemblies are fragmented the possibility of erroneously circularizing non-circular contigs arises, hereon termed false circularization. False circularization is of particular concern because one cause of assembly fragmentation is the presence of large repeat sequences in the assembled DNA. Contig breaks are often associated with these repeats, and therefore the presence of overlapping sequence at both ends of a contig does not necessitate that the contig is circular. Such a situation is particularly problematic for overlap approaches to circularization. Indeed, the majority of false circularizations made by the methods in our comparison occurred in one sample, the *Bacillus subtilis* strain NCTC3610, whose assembly was the most fragmented of our test panel. For this sample, Minimus2 falsely circularized two fragments of the chromosome, and BLAST falsely circularized 5 contigs including 4 small contigs representing repeat sequences and one fragment of the chromosome. The only false circularization made by Circlator was also in NCTC3610, where it circularized a small contig representing a repeat sequence. In total, Minimus2 falsely circularized 3 contigs, and BLAST falsely circularized 7.

Assembly of long reads without accounting for circularization can provide particularly problematic results for small plasmids whose length is shorter than the length of the reads used to assemble it. This can lead to the production of contigs containing the entire sequence of the plasmid two or more times (Figure 1c). Circlator and our iterative BLAST approach correctly identified these occurrences and collapsed the plasmid sequences down to a single copy, whereas Minimus2 recognized only the first copy of the repeat sequence at each end of the contig, leading to the creation of an incorrectly circularized contig in some cases (for example the plasmids of sample NCTC13360, as shown in Supplementary Figure S16).

#### Polishing

Quiver was run on the output of HGAP, and on the output of each of the circularization programs. All assemblies, both pre- and post-Quiver, were evaluated using QUAST [25] version 2.3 with the options `--gage –R` to use a reference sequence and run the GAGE [26] analysis. Complete QUAST results are given in Supplementary table S3. Summary plots are shown in Supplementary Figure S23, where it can be seen that, after running Quiver, Circlator generally produces higher quality assemblies than the other approaches. These plots also highlight that assembly polishing using Quiver is critical, regardless of which circularization method is used.

### *P. falciparum* apicoplast and mitochondrion

Another test set is the *P. falciparum* genome that consists of 14 linear chromosomes plus circular mitochondrion and apicoplast sequences. We evaluated the circularization tools using data from the reference strain 3D7 [27]. Since the latest available version of the apicoplast has a large deletion caused by a near-identical inverted repeat, we generated an improved apicoplast sequence to use as a reference sequence when determining the accuracy of the circularization tools (for details see Supplementary Material).

Reads from 11 PacBio SMRT cells (accessions ERR951787 to ERR951797 inclusive) were assembled using HGAP. Contigs matching the existing apicoplast and mitochondrion reference sequences were identified by taking all hits reported by nucmer (with default settings for nucmer, and `delta-filter -i 95 -l 1000`). The resulting two contigs, one for each of the apicoplast and mitochondrion, were input to the Minimus2 and BLAST methods. Circlator was run using the same two contigs, together with all corrected reads that mapped to those contigs using BWA MEM with the option `-x pacbio.` Since the reads were low coverage, the kmer used by SPAdes was set to 101 (with Circlator option `--assemble_spades_k 101`) in order for it to output complete assemblies. All three tools correctly circularized the apicoplast. Figure 3 shows a comparison of the HGAP assembly, which comprises many copies of the apicoplast sequence, against the output of circlator. Minimus2 and BLAST attempted to circularize the mitochondrion, but failed to recognise the multiple copies in the input contig, and output several copies of the sequence. Of the three tools, only Circlator correctly circularized the mitochondrion sequence (Figure Supplementary Figure S21).

**Figure 3:**
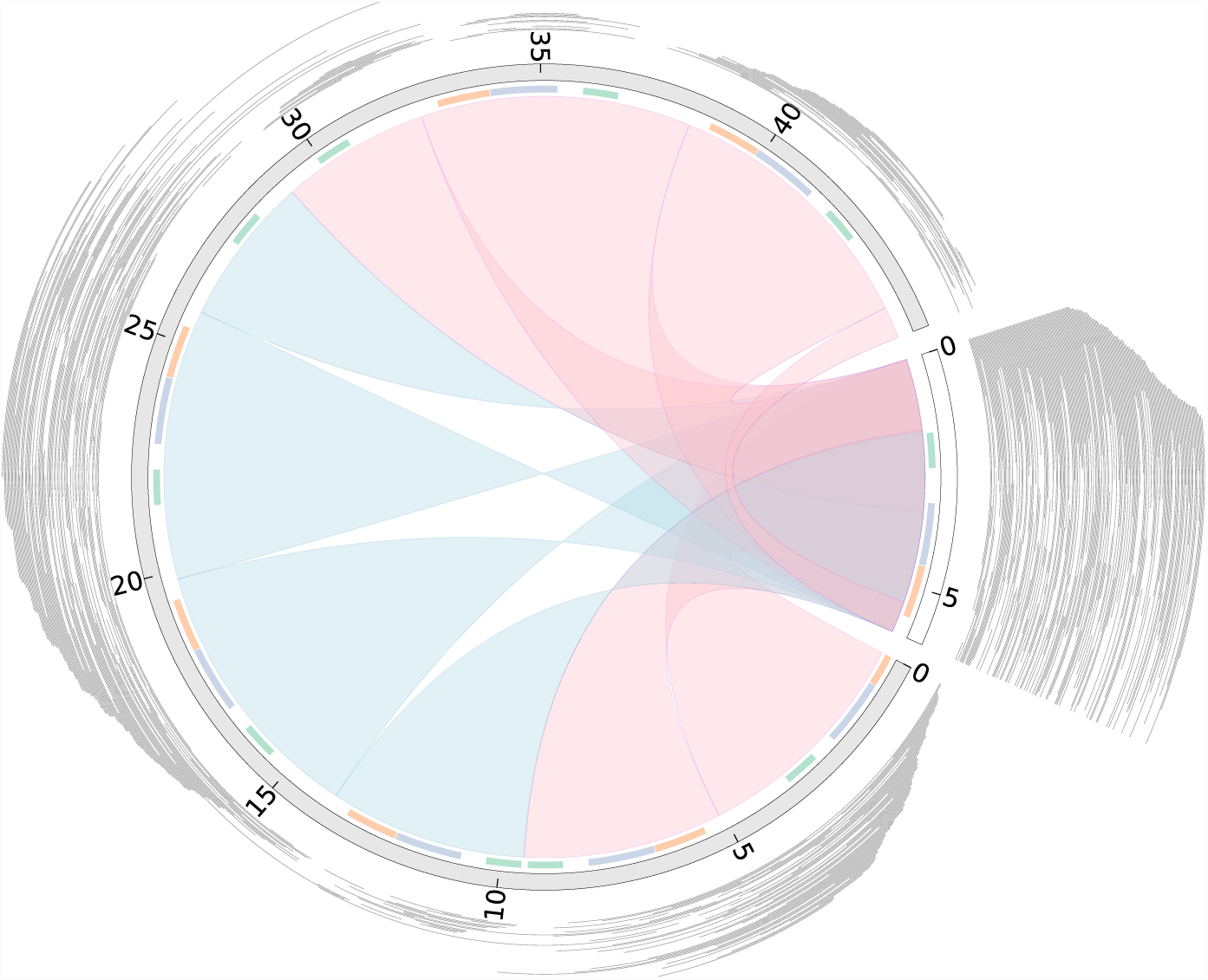
Comparison of HGAP assembly of *P. falciparum* apicoplast and Circlator output. The HGAP and Circlator assemblies are shown in gray and white respectively, with the numbers showing lengths in kilobases. Nucmer matches between the genomes are shown as blue (hits in the same orientation) and pink (hits in opposing orientations). Matches to the three apicoplast genes, cox1 (blue), cox3 (green) and cob (orange), are shown as a coloured track inside the assemblies. The corrected reads mapped to each of the assemblies are shown in grey outside the assemblies. This figure was generated using Circos [28].

### *H. sapiens* mitochondrion

To determine the applicability of circularization to the human mitochondrial genome, a test assembly and set of corrected PacBio reads was generated from a shotgun sequence dataset (accession numbers SRR1304331 to SRR1304530 inclusive). To extract just the reads that correspond to the mitochondrion genome, first the reads were mapped to the reference GRCh38 mitochondrion sequence (NC_012920.1) using BWA MEM with the option `-x pacbio.` To remove false-positives, the reads that mapped were then mapped to the entire GRCh38 genome using BWA MEM with the same settings. Only reads with a primary match to the mitochondrion were retained, and assembled with PBcR version 8.3rc2 using the options `–maxCoverage 1000 -length 500 -partitions 200 genomeSize=16569`. PBcR output a single contig, containing more than two copies of the mitochondrion sequence. Minimus2 attempted to circularize the contig but produced errors, whereas BLAST and Circlator correctly circularized this contig.

### *E. coli* Nanopore data

We explored the possibility of circularizing a nanopore assembly, which is more challenging than PacBio because of the higher error rate in the reads and assembly contigs, using *Escherichia coli* ONT MinION data [22]. We ran a *de novo* assembly of the reads using PBcR, in order to generate the contigs and corrected reads required as input to Circlator, using the data and instructions at [29] and checkout revision 4642 of the source code. Since the assembly and corrected reads are of lower quality than those of PacBio, it was required to make Circlator more permissive when matching the input assembly contigs to the SPAdes contigs generated by Circlator using the options `--merge_min_id 85 --merge_breaklen 1000`. These change the default MUMmer parameters of Circlator, which are tuned to PacBio data, by passing -b 1000 to nucmer and -i 85 to delta-filter instead of the default `-b 500` and `-i 95`. This lowers the minimum percent identity from 95 to 85 (-i 85) and helps to extend hits through poorly aligned regions (`-b 1000`). Minimus2 and BLAST failed to circularize the assembly, however, Circlator successfully produced a correctly circularized genome sequence.

## Conclusions

Here we have presented the first tool that automatically resolves circular genome assemblies. Since circularization was the only remaining stage of genome assembly that required manual work, Circlator completes the automation of the process of assembling raw reads into a finished genome sequence. It was successfully applied to a wide range of species and different technologies and outperformed existing semiautomatic methods. In conclusion, Circlator provides the final step in the automated production of reference quality long read genome assemblies.

## Software

Circlator is open source and available for Linux at http://sanger-pathogens.github.io/circlator/ under the GPLv3 licence. It has low memory usage and a short run time (see Supplementary text, Supplementary Tables S4,S5 and Supplementary Figure S24). Circlator is easy to use, with a single call required to run the whole pipeline, and is also modular, so that any stage of the pipeline can be run in isolation.

## List of abbreviations

NCTC: National Collection of Type Cultures
ONT: Oxford Nanopore Technologies
PacBio: Pacific Biosciences

## Author’s contributions

MH and SH conceived the project and Circlator algorithm. MH and NDS wrote the software. MH and SH ran and analysed the benchmarking of the tools. TO produced the *P. falciparum* reference apicoplast sequence. All authors wrote, read and approved the final manuscript.

## Acknowledgements

We thank Matthew Berriman and Chris Newbold for permission to use the *P. falciparum* sequencing data and Mandy Sanders for her contribution on the *P. falciparum* apicoplast. This work was supported by the Wellcome Trust grant [098051].

## Additional Files

### Additional file 1 — supplementary_material.pdf

Supplementary text and figures supporting the main text.

### Additional file 2 — supplementary_tables.xls

An Excel spreadsheet containing Supplementary Tables S1–4.

Supplementary table S1: NCTC project IDs and accession numbers for the PacBio bacterial samples.

Supplementary table S2: A complete breakdown of results on the PacBio bacterial samples (an expanded version of Table 1).

Supplementary table S3: QUAST results on the PacBio bacterial samples. Supplementary table S4: Run time and memory usage for all data sets.

### Additional file 3 — ftp://ftp.sanger.ac.uk/pub/pathogens/circlator/Supplementary_data/Circlator_supplementary_data.tar.gz

A gzipped archive containing all scripts, corrected reads, assemblies and other data necessary to reproduce the results.

